# A two-lane mechanism for selective biological ammonium transport

**DOI:** 10.1101/849562

**Authors:** Gordon Williamson, Giulia Tamburrino, Gaëtan Dias Mirandela, Mélanie Boeckstaens, Marcus Bage, Andrei Pisliakov, Callum M. Ives, Eilidh Terras, Adriana Bizior, Paul A. Hoskisson, Anna Maria Marini, Ulrich Zachariae, Arnaud Javelle

## Abstract

The transport of charged molecules across biological membranes faces the dual problem of accommodating charges in a highly hydrophobic environment while maintaining selective substrate translocation. A particular controversy has existed around the mechanism of ammonium exchange by the ubiquitous Amt/Mep/Rh transporter family, an essential process in all kingdoms of life. Here, using a combination of electrophysiology, yeast functional complementation and extended molecular dynamics simulations, we reveal a unique two-lane pathway for electrogenic NH_4_^+^ transport in two archetypal members of the family. The pathway underpins a mechanism by which charged H^+^ and neutral NH3 are carried separately across the membrane after NH_4_^+^ deprotonation. This mechanism defines a new principle of achieving transport selectivity against competing ions in a biological transport process.

## Introduction

The transport of ammonium across cell membranes is a fundamental biological process in all kingdoms of life. Ammonium exchange is mediated by the ubiquitous ammonium transporter/methylammonium permease/Rhesus (Amt/Mep/Rh) protein family. The major role of bacterial, fungal, and plant Amt/Mep proteins is to scavenge ammonium for biosynthetic assimilation, whereas mammals are thought to produce Rh proteins in erythrocytes, kidney, and liver cells for detoxification purposes and to maintain pH homeostasis (*1, 2*). In humans, Rh mutations are linked to pathologies that include inherited hemolytic anemia, stomatocytosis, and early-onset depressive disorder (*2*). Despite this key general and biomedical importance, so far, no consensus on the pathway and mechanism of biological ammonium transport has been reached.

High resolution structures available for several Amt and Rh proteins show a strongly hydrophobic pore leading toward the cytoplasm (*3-6*). This observation had led to the conclusion that the species translocated through Amt/Mep/Rh proteins is neutral NH_3._ However, this view has recently been experimentally challenged, first for some plant Amt proteins (*7-10*), followed by further *in-vitro* studies revealing that the activity of bacterial Amt proteins is electrogenic (*11, 12*). Computational (*13*) and experimental studies (*14*) further demonstrated that deprotonation of NH_4_^+^ is likely to be a major step in ammonium transport. Taken together, these findings renewed a long-standing debate on the mechanism by which a charge is translocated through a hydrophobic pore and how selectivity for NH_4_^+^ over competing ions is achieved.

Here, we reveal the pathways, mechanism, and key determinants of selectivity of electrogenic ammonium transport in Amt and Rh proteins, unifying the diverse observations that led to these seemingly incompatible suggestions. The transport mechanism is underpinned by the separate transfer of H^+^ and NH_3_ on a unique two-lane pathway following NH_4_^+^ sequestration. This mechanism ensures that ammonium – which occurs mainly in protonated form in the aqueous phase – is efficiently translocated across the membrane, while maintaining strict selectivity against K^+^, a monovalent cation of similar size. This previously unobserved principle is likely to form a new paradigm for the electrogenic members of the Amt/Mep/Rh family. Similar mechanisms may be utilized by other membrane transporters to facilitate the selective translocation of pH-sensitive molecules.

## Results and Discussion

Motivated by our finding that the activity of *Escherichia coli* AmtB is electrogenic (*12*), we first investigated the Rh50 protein from *Nitrosomonas europaea* (NeRh50). Rh and Amt proteins are distant homologs, and thus a functional distinction between both subfamilies was proposed (*2*). The architecture of NeRh50 is highly representative of Rh proteins (*4, 5*) which have been reported to serve as electroneutral NH_3_ or CO_2_ gas channels (*5, 15-18*). The activity of purified NeRh50 reconstituted into liposomes was quantified using Solid-Supported Membrane Electrophysiology (SSME) (*19, 20*), where we recorded a NH_4_^+^-selective current (Fig. 1) with a decay rate that is strongly dependent on lipid-to-protein ratio (LPR; Table S4). Furthermore, expressed in a *Saccharomyces cerevisiae* triple-*mepΔ* strain, which is deprived of its three endogenous Mep ammonium transporters, NeRh50 complemented the growth defect on minimal medium containing ammonium as sole nitrogen source (Fig. 1).

**Fig. 1:**
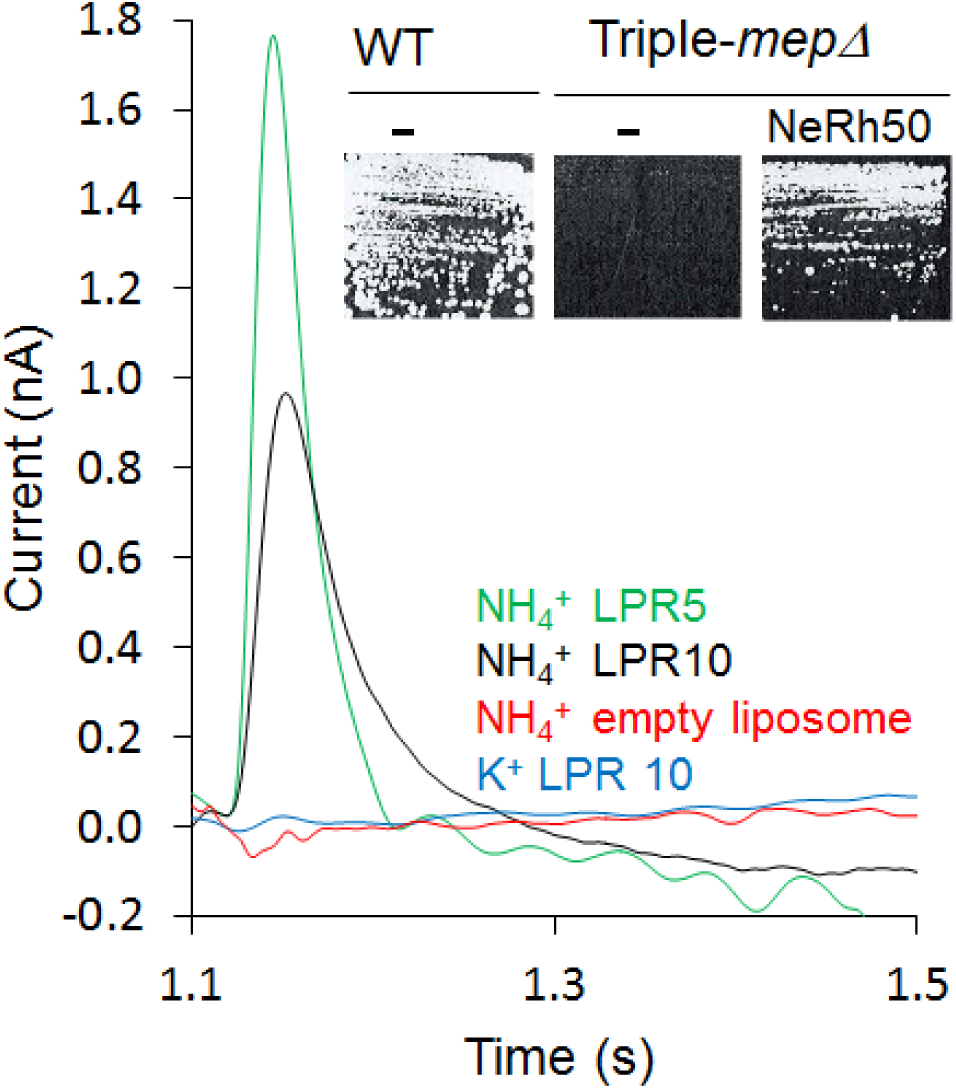
Characterization of the activity of NeRh50. SSME analysis. Transient current measured on proteoliposomes after a 200 mM pulse (NH_4_^+^ or K^+^). ***Insert:*** Yeast complementation by NeRh50. Growth test on minimal medium supplemented with 3 mM ammonium as sole nitrogen source.

The electrogenic transport activity and NH_4_^+^ selectivity we observed for NeRh50 and AmtB suggested a conserved transport mechanism amongst the distant Amt and Rh proteins and further highlighted the question of how these proteins achieve selective charge translocation through their hydrophobic pore. Since the most substantive body of structural information is available for the paradigmatic ammonium transporter, AmtB from *E. coli* and its variants, we next aimed to decipher the molecular mechanism of electrogenic NH_4_^+^ transport in AmtB. AmtB forms homotrimers in the cytoplasmic membrane, in which each monomer exhibits a potential periplasmic NH_4_^+^ binding region (S1, near residues S219 and D160) followed by a strongly hydrophobic pore leading toward the cytoplasm (Fig. 2) (*3*). Two highly-conserved histidine residues, H168 and H318, protrude into the lumen, forming the family’s characteristic “twin-His” motif (Fig. 2) (*21*). Previous computational studies suggested that NH_4_^+^ is deprotonated after binding near S1 via residues S219 or D160, with subsequent return of H^+^ to the periplasm (*22-24*). As our experiments demonstrated that AmtB activity is electrogenic, we excluded return of the proton into the periplasm after NH_4_^+^ deprotonation.

**Fig. 2:**
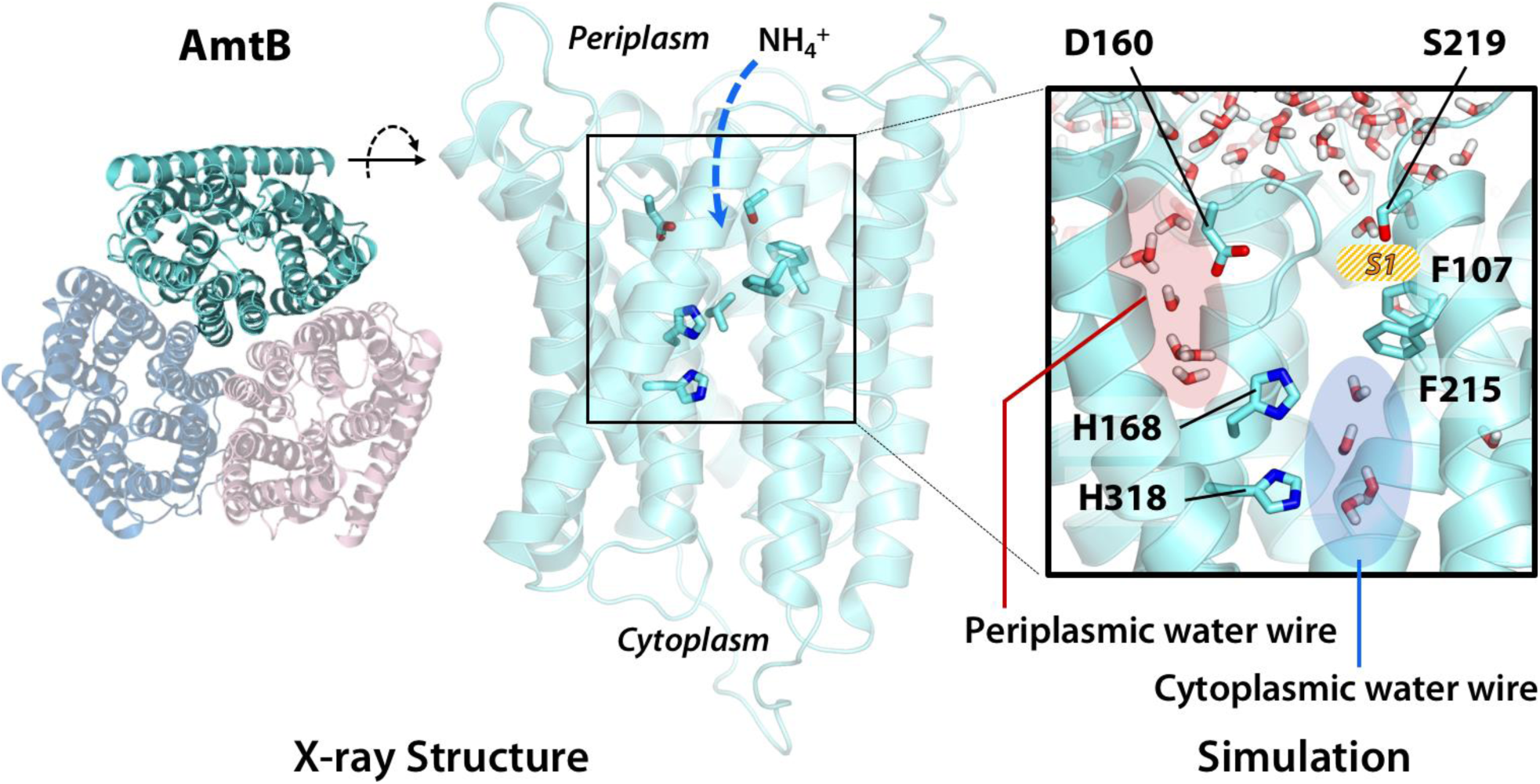
Formation of the two water wires in AmtB. Extended atomistic simulations show a hydration pattern across the protein, in which cytoplasmic and periplasmic water wires, connected via H168, form a continuous pathway for proton transfer from the S1 NH_4_^+^ sequestration region to the cytoplasm.

To investigate the role of residues S219 and D160 in the context of electrogenic transport, we expressed wild-type (wt) AmtB and the variants S219A and D160A in *S. cerevisiae* triple-*mepΔ.* Using ammonium as the sole nitrogen source, we found equivalent growth of cells expressing AmtB^S219A^ or AmtB, while cells expressing AmtB^D160A^ failed to grow, showing that AmtB^D160A^ is unable to replace the function of the endogenous Mep transporters (Fig. 3A). The activity of purified AmtB variants reconstituted into liposomes was quantified using SSME. Electrogenic transport activity, triggered by a 200 mM ammonium pulse, led to a transient current with a maximum amplitude of 3.38 nA in AmtB, while AmtB^S219A^ and AmtB^D160A^ displayed reduced maximum currents of 1.86 nA and 0.63 nA, respectively (Fig. 3A, Fig. S3A). In contrast to AmtB^S219A^ however, the lifetime of the AmtB^D160A^ current was unaffected by changes in liposomal LPR (Table S4, Fig. S3B), and the catalytic constants (*K*_*m*_) for AmtB and AmtB^S219A^ were identical, while the *K*_*m*_ of AmtB^D160A^ was increased by ∼70-fold (Fig. 3A). Taken together, these results show that the AmtB^S219A^ variant sustains the complete transport cycle, albeit with a slightly lowered transport rate due to the disruption of the S1 region. In contrast, AmtB^D160A^ is transport-deficient and the small transient current arises from the binding of a NH_4_^+^ to the protein (*20*). Accordingly, our results rule out an essential function of S219 in NH_4_^+^ deprotonation, while they demonstrate a key role of the highly-conserved D160 in the transport activity. In accordance with previous computational (*23, 25*) and experimental studies (*26, 27*), we therefore concluded that D160 is involved in NH_4_^+^ deprotonation. However, in contrast to these previous studies, we hypothesized that the proton is subsequently transferred to the cytoplasm.

**Fig. 3:**
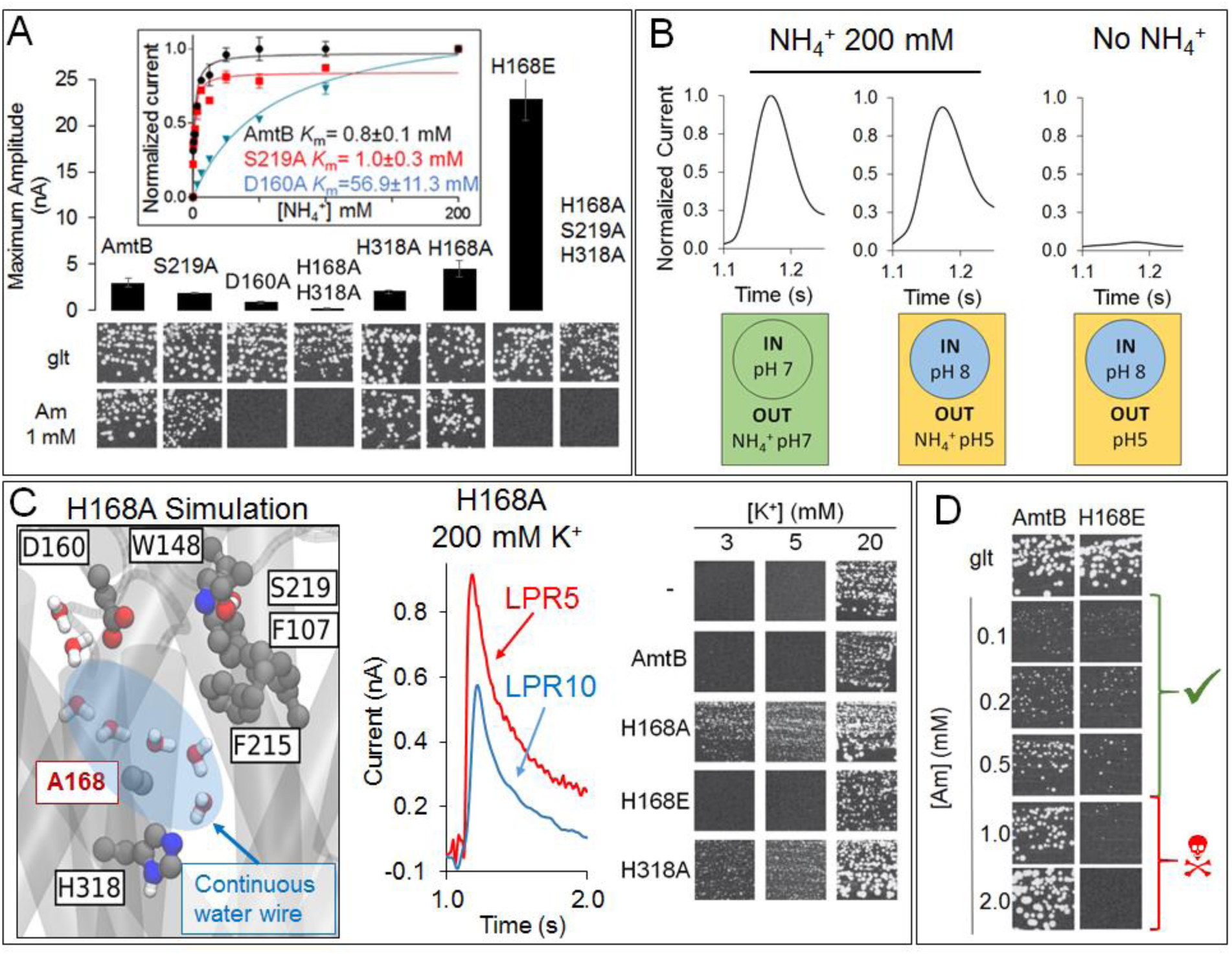
Characterization of AmtB translocation pathways. (A) *In vivo* and *in vitro* analysis of AmtB variants affected in the translocation pathways. *Upper panel:* maximum amplitude of the transient current measure using SSME. *Lower panel*: yeast complementation test. Glutamate (glt) or 1 mM ammonium as a sole nitrogen source. *Insert*: Kinetics analysis. The maximum amplitudes recorded after a 200 mM ammonium pulse have been normalised to 1.0 for comparison. (B) Effect of a proton gradient on AmtB activity. The transient currents were measured following an ammonium pulse at pH 7 (left) or under an inwardly directed pH gradient in the presence (centre) or absence (right) of ammonium. (C) The H168A and H318A variant loses NH_4_^+^ specificity. *Left*: MD simulation of AmtB^H168A^ showing formation of a continuous water wire traversing the central pore region. *Centre:* Transient current measured after a potassium pulse in proteoliposomes containing AmtB^H168A^. *Right*: Growth tests of *Saccharomyces cerevisiae* triple-*mepΔ trk1Δ trk2Δ* yeast cells on solid minimal medium containing different KCl concentrations. (D) The H168E variant is toxic when expressed in yeast. Yeast complementation test of wt AmtB compared to AmtB^H168E^. No complementation (i.e. toxicity) was observed for AmtB^H168E^ in medium containing more than 1 mM ammonium.

We then aimed to identify dynamic polar networks across AmtB that could form a transfer pathway for H^+^ from the NH_4_^+^ deprotonation site to the cytoplasm. In the absence of NH_4_^+^, a proton pulse did not trigger a discernible current and additionally, in the presence of ammonium, an inward-orientated pH gradient did not increase AmtB activity (Fig. 3B). These findings suggested that the current associated with AmtB activity is generated by specific deprotonation rather than an H^+^-dependent symport activity. To locate potential polar transfer routes, we performed atomistic molecular dynamics (MD) simulations of AmtB in mixed lipid bilayers, which initially showed the hydration of a portion of the putative hydrophobic NH_3_ pathway, spanning from the twin-His motif to the cytoplasm (cytoplasmic water wire – CWW; Figs. 2, S4). The presence and occupancy of the CWW confirmed previous observations (*28*). Importantly however, over longer timescales, our simulations revealed the presence of a previously unidentified second water-filled channel (periplasmic water wire - PWW), which opens from residue D160 to the central twin-His motif (Figs. 2, S5). The PWW is not formed when D160 is mutated to alanine, in line with our data showing that the AmtB^D160A^ mutant is inactive (Fig. 3A, S6). The CWW and PWW are connected via the twin-His motif, which bridges the aqueous chains, while preventing the formation of a continuous water channel in the simulations (Fig. 2). Although neither the CWW nor the PWW are sufficiently wide to allow the transfer of solvated NH_4_^+^, water molecules and histidine side chains often serve as efficient pathways to facilitate proton transfer in proteins (*29*). Further calculations suggested that H^+^ transfer between the water molecules in the PWW and CWW is possible and occurs with high rates (the highest energy barrier is ∼18 kJ/mol in the cytoplasmic wire near the twin-His motif; Table S5).

We also probed if the twin-His motif enables proton transfer between the two water wires by recording the activities of twin-His variants. Earlier calculations showed that H^+^ transfer within the twin-His motif parallel to the CWW is possible (*13*). Expressed in *S. cerevisiae* triple-*mepΔ*, we found that AmtB^H168A/H318A^ did not support cell growth (Fig. 3A). *In-vitro* SSME measurements displayed LPR-independent current decay rates (Fig. S3C, Table S4), showing that the residual current was caused by the association of NH_4_^+^ to AmtB without further transport (*20*). No current was recorded for the triple mutant AmtB^S219A/H168A/H318A^, in which binding at the periplasmic face was additionally altered, confirming that the residual current reflects NH_4_^+^ interaction near S1 (Fig. 3A, Fig. S3C). The double-His mutant AmtB^H168A/H318A^ is thus able to bind NH_4_^+^, but is no longer capable of transporting the substrate across the membrane, in line with our previous structural analysis which showed the CWW to be absent in the pore of the double-His mutant AmtB^H168A/H318A^ (*21*). In the single His variant AmtB^H168A^ however, we unexpectedly find a LPR dependent current in our SSME recordings (Fig. 3A, Fig. S3C, Table S4). Furthermore, triple-*mepΔ* yeast cells expressing AmtB^H168A^ display growth in the presence of low ammonium concentrations (Fig. 3A). Our previous crystal structure (*21*) and our MD simulations (Fig. 3C) show increased hydration in the area around A168, which could potentially form a pathway for efficient H^+^ transfer or direct translocation of NH_4_^+^, thereby explaining the growth on NH_4_^+^.

This data suggested that the twin-His motif is not essential for NH_4_^+^ transport despite its conservation. Therefore, we repeated our SSME experiments on AmtB^H168A^ using the competing K^+^ ion as substrate, since earlier studies implicated a role of the twin-His motif in AmtB selectivity (*30*). A 200 mM K^+^ pulse triggered a current in AmtB^H168A^ whose decay rate strongly depended on LPR (Fig. 3C, Table S4). Growth tests of a *S. cerevisiae* strain lacking its three endogenous Mep (NH_4_^+^) and two Trk (K^+^) transporters in minimal medium containing a limited concentration of K^+^ showed that the variant AmtB^H168A^, but not wt-AmtB complemented the growth defect (Fig. 3C). We obtained similar results for the variant AmtB^H318A^, where growth complementation in both NH_4_^+^ and K^+^ containing minimal medium was observed (Fig. 3A, C).

Taken together, these results demonstrate that both AmtB^H168A^ and AmtB^H318A^ translocate K^+^ ions across the membrane, and that these substitutions within the twin-His motif thus abolished selectivity for NH_4_^+^. Therefore, the presence of both histidine residues is critical, since permeability of ammonium transporters for K^+^ would compromise ionic homeostasis and disrupt the membrane potential of *E. coli* cells, which crucially depends on maintaining K^+^ concentration gradients across the membrane. Moreover, since AmtB is expressed in *E. coli* under nitrogen starvation conditions (low NH_4_^+^/K^+^ ratio), the loss of selectivity for NH_4_^+^ would impede ammonium uptake. Our results thus demonstrate that the twin-His motif, which is highly conserved amongst members of the family, is an essential functional element in the transport mechanism, preventing the transport of competing cations, whilst providing a pathway for proton transfer by bridging the periplasmic and the cytoplasmic water wires.

Despite high conservation in the Mep/Amt/Rh family, a substitution of H168 to E is found in some fungal Mep proteins (*31*). In our SSME recordings, we found the AmtB variant H168E to be super-active with an 8-fold increased current compared to wild-type (Fig. 3A, Fig. S3C), and a strongly LPR-dependent decay rate (Table S4). In this variant, NH_4_^+^ selectivity is retained as K^+^ pulses do not trigger any measurable current (Fig. 3C and Table S4). Expressed in the yeast triple-*mepΔ*, however, we found that the super-active phenotype is cytotoxic (Fig. 3D). Together with our previous crystal data on the internal hydration of AmtB^H168E^ (Fig. S8), these findings suggest that the additional charge introduced by the H168E substitution further stabilizes the periplasmic and cytoplasmic water wires, raising proton transfer rates, while the large glutamate side chain does not allow the formation of a direct connection between the water chains. The drastically increased activity of this variant is, however, not tolerated by all cell types.

Additional free-energy calculations we performed showed that NH_3_ permeation through the hydrophobic pore of AmtB experiences a maximum energy barrier of ∼10 kJ/mol (Fig. S9). Since both energy barriers for H^+^ transfer along the water chains and NH_3_ permeation are relatively small, either initial deprotonation or proton transfer across the twin-His motif could be rate-limiting for overall NH_4_^+^ transport. Since we found a strongly increased activity for the H168E variant, we conclude that the transport rate of AmtB is determined by the rate of H^+^ transfer across the twin-His motif.

A new model for the mechanism of electrogenic ammonium transport therefore emerges from our findings (Fig. 4). After deprotonation of NH_4_^+^ at the periplasmic side, a previously undiscovered polar conduction route enables H^+^ transfer into the cytoplasm. A parallel pathway, lined by hydrophobic groups within the protein core, facilitates the simultaneous transfer of uncharged NH_3_, driven by concentration differences. On the cytoplasmic face, the pH of the cell interior leads to recombination to NH_4_^+^, most likely near a second hydrophobic gate (Fig. S9). The major kinetic barrier to transport occurs for H^+^ transfer across the twin-His motif, which bridges the water chains and thereby constitutes the major selectivity gate for NH_4_^+^ transport. We propose that this mechanism is conserved amongst the electrogenic members of the Amt/Mep/Rh family.

**Fig. 4:**
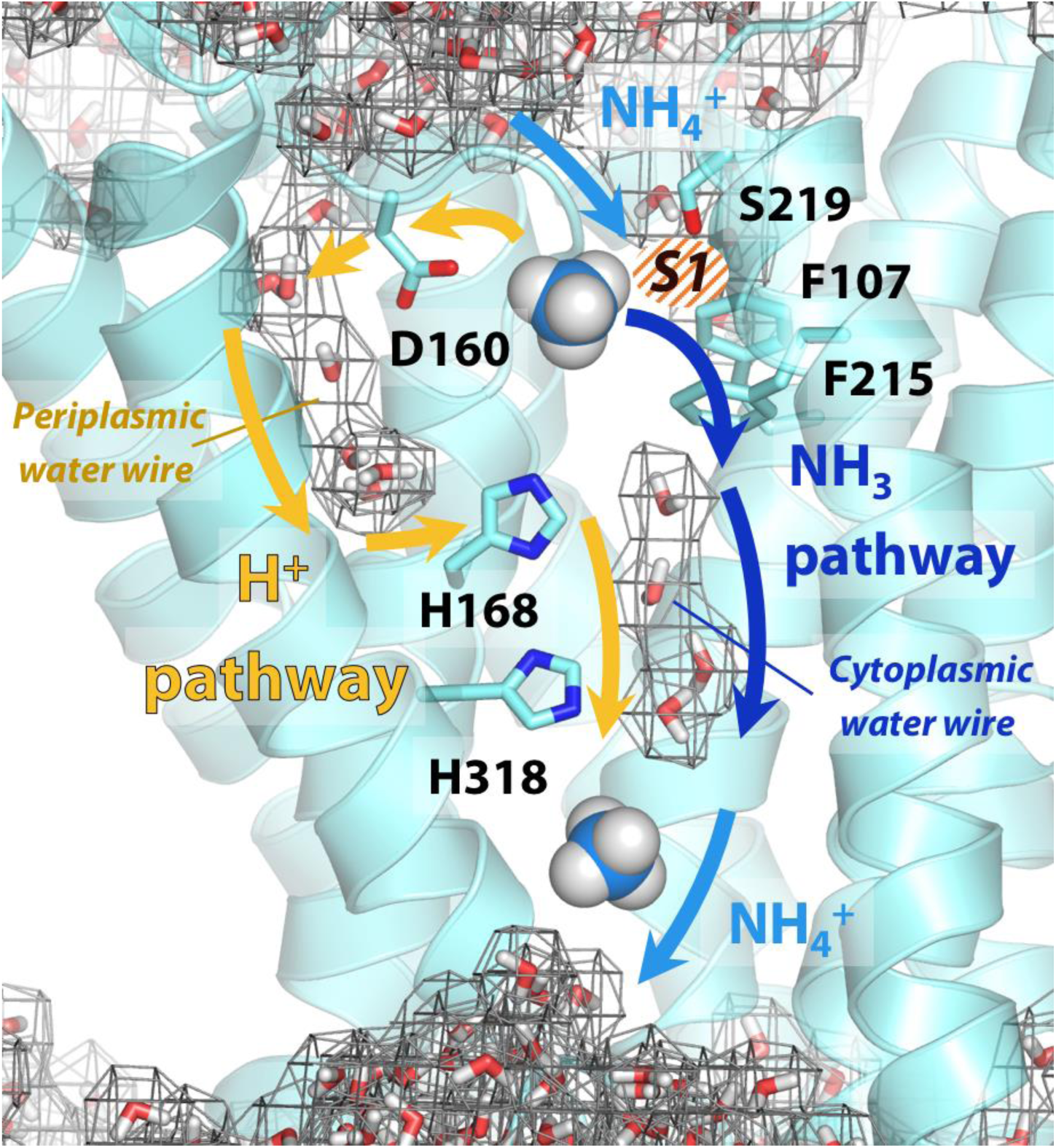
Mechanism of electrogenic NH_**4**_^**+**^ translocation in AmtB. Following sequestration of NH_4_^+^ at the periplasmic face, NH_4_^+^ is deprotonated and H^+^ and NH_3_ follow two separated pathways to the cytoplasm (orange arrows depict the pathway for H^+^ transfer, dark blue arrows for NH_3_), facilitated by the presence of two internal water wires. Recombination likely occurs near the cytoplasmic exit (Fig. S9). The hydrated regions within the protein as observed in simulations are highlighted by wireframe representation, crucial residues involved in the transport mechanism are shown as sticks.

## Conclusion

Our findings define a new mechanism, by which ionizable molecules that are usually charged in solution are selectively and efficiently transported across a highly hydrophobic environment like the AmtB/Rh pore. Alongside size-exclusion and ion desolvation (*32*), it adds a new principle by which selectivity against competing ions can be achieved. Other biological transport systems, like the formate/nitrite transporters, may share similar mechanisms involving deprotonation-reprotonation cycles (*33*).

## Materials and Methods

### Mutagenesis

AmtB mutants were generated using the Quikchange XL site-directed mutagenesis kit (Agilent Technologies), according to the manufacturer’s instructions. The primers used for mutagenesis are listed in Table S1. The template was the *amtB* gene cloned into the plasmid pET22b(+), as previously described (*34*)(Table S2).

#### AmtB and NeRh50 expression in yeast and complementation test

The different variants of *amtB* were amplified using *amtB* cloned into pET22b(+) (Table S2) as a template with the primers AmtB XhoI and AmtB BamHI (Table S1) and then sub-cloned into the plasmids pDR195 (Table S2). The NeRh50 gene was amplified from *N. europaea* genomic DNA (kind gift from Daniel J. Arp and Norman G. Hommes, Department of Botany and Plant Pathology, Oregon State University, Corvallis, USA) using the primers P5’NeRh and P3’NeRh (Table S1), and was then cloned into the SpeI and EcoRI restriction sites of the high-copy vector p426Met25 (Table S2), allowing controlled-expression of NeRh50 by the yeast methionine repressible MET25 promoter.

### Yeast growth assays

*Saccharomyces cerevisiae* strains used in this study are the 31019b strain (*mep1Δ mep2Δ mep3Δ ura3*) and the 228 strain (*mep1Δ mep2Δ mep3Δ trk1Δ trk2Δ leu2) (31, 35)*. The plasmids used in this study are listed in Table S1. Cell transformation was performed as described previously (*36*). For growth tests on limiting ammonium concentrations, yeast cells were grown in minimal buffered (pH 6.1) medium and for growth tests on limiting potassium concentrations, a minimal buffered (pH 6.1) medium deprived of potassium salts was used (*37*). 3% glucose was used as the carbon source and, 0.1% glutamate, 0.1% glutamine or (NH_4_)_2_SO_4_ at the specified concentrations were used as the nitrogen sources.

### Protein purification

AmtB(His_6_) cloned into the pET22b(+) vector (Table S2) was overexpressed and purified as described previously (*34*). The plasmid pAD7 (Table S2) was used to overexpress NeRh50 in the *E. coli* strain GT1000 (*26*). GT1000 was transformed with pAD7 and grown in M9 medium (*38*), in which ammonium was replaced by 200 μg/ml glutamine as sole nitrogen source. NeRh50 was purified as described by Lupo *et al.* (*5*) with minor modifications, namely: the membrane was solubilised using 2% lauryldecylamine oxide (LDAO) instead of 5% *n*-octyl-β-D-glucopyranoside (OG), and 0.09% LDAO was used in place of 0.5% β-OG in the final size exclusion chromatography buffer (50 mL Tris pH 7.8, 100 mL NaCl, 0.09% LDAO).

#### AmtB and NeRh50 insertion into proteoliposomes

AmtB and NeRh50 were inserted into liposomes containing *E. coli* polar lipids/phosphatidylcholine (POPC) 2/1(wt/wt) as previously described (*12*). For each AmtB variant, proteoliposomes were prepared at lipid-to-protein ratios (LPRs) of 5, 10, and 50 (wt/wt). The size distribution of proteoliposomes was measured by dynamic light scattering (DLS) using a Zetasizer Nano ZS (Malvern Instruments). This analysis showed that the proteoliposomes had an average diameter of 110 nm (Fig. S1). Proteoliposomes were divided into 100 µL aliquots and stored at −80°C.

The quantity of protein inserted in liposomes was assessed by SDS-PAGE analysis (Fig. S2A). 15 µL of proteoliposomes (5mg/ml) containing each variant at a LPR of 5:1, 10:1, or 50:1 (w/w), or wild-type AmtB at LPR 10, were run on 10% SDS-PAGE gels, demonstrating that the quantity of protein inserted is similar for all variants according to the respective LPR. It was noted that AmtB^D160A^ ran as a monomer in SDS-PAGE gel, rather than the trimer seen for other variants. To ensure that all AmtB variants were correctly inserted into the proteoliposomes, the proteoliposomes were solubilised in 2% DDM and the proteins analyzed by size exclusion chromatography using a superdex 200 (10×300) enhanced column. The elution profile of all variants and the wild-type were identical, showing a single monodisperse peak eluting between 10.4-10.6 ml. This demonstrated that all proteins were correctly folded, as trimers, in the proteoliposomes (Fig. S2B).

### Solid Supported Membrane Electrophysiology

To form the solid-supported membrane, 3 mm gold-plated sensors were prepared according to the manufacturer’s instructions (Nanion Technologies), as described previously (*19*). Proteoliposomes/empty liposomes were defrosted and sonicated in a sonication bath at 35 W for 1 min, diluted 10 times in non-activating (NA) solution (Table S3), and then 10 µl were added at the surface of the SSM on the sensor. After centrifugation, the sensors were stored at 4°C for a maximum of 48 h before electrophysiological measurements.

All measurements were made at room temperature (21°C) using a SURFE^2^R N1 apparatus (Nanion Technologies) with default parameters (*19*). Prior to any measurements, the quality of the sensors was determined by measuring their capacitance (15-30 nF) and conductance (<5 nS).

For functional measurements at a fixed pH, a single solution exchange protocol was used with each phase lasting 1 second (*19*). First, non-active (NA) solution was injected onto the sensor, followed by activating (A) solution containing the substrate at the desired concentration and finally NA solution (Table S3).

For the measurements under inwardly orientated pH gradient, a double solution exchange protocol was used (*19*), in which an additional resting solution phase of 15 min in NA solution at pH 8 was added to the end. The incubation phase adjusts the inner pH of the proteoliposomes to pH 8 and establishes a pH gradient at the beginning of each measurement by pulsing the activation solution at pH 5.

The activity of AmtB and NeRh50 was tested at pH 7 before and after each measurement to ensure that there was no activity loss during the measurements. All measurements were recorded on 6 sensors from two independent protein purification batches, with 3 measurements recorded for each sensor.

The kinetic parameters were calculated using Graphpad Prism 6 and fitted according to the Michaelis-Menten equation. Lifetime analysis of the current (decay time of the transient current) was performed to differentiate small pre-steady state currents, which arise due to the binding of a charged species to membrane proteins, from currents reflecting full transport cycles, which show faster decay rates under raised liposomal LPR (*19*). The decay time of the transient current (Table S4) was calculated by fitting the raw transient current data between the apex of the peak and the baseline (after transport) with a non-linear regression using OriginPro 2017 (OriginLab). The regression was done using a one-phase exponential decay function with time constant parameter:

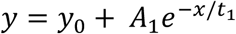

The fit was obtained using the Levenberg-Marquardt iteration algorithm, where *x* and *y* represent coordinates on the respective axis, *y*_*0*_ represents the offset at a given point, *A* represents the amplitude, and *t* is the time constant.

### Molecular Dynamics Simulations

The AmtB trimer (PDB code: 1U7G) (*3*) was processed using the CHARMM-GUI web server (*39*). Any mutations inserted during the crystallization process were reverted to the wild-type form. The N-termini and C-termini of the subunits were capped with acetyl and N-methyl amide moieties, respectively. The protein was then inserted into a membrane patch of *xy*-dimensions 13×13 nm. Unless otherwise specified, a membrane composition of palmitoyl oleoyl phosphatidyl ethanolamine and palmitoyl oleoyl phosphatidyl glycine (POPE/POPG) at a 3:1 ratio was used in order to approximate the composition of a bacterial cytoplasmic membrane. We employed the CHARMM36 forcefield for the protein and counter ions (*40*). The water molecules were modeled with the TIP3P model (*41*). Water bonds and distances were constrained by the Settle method (*42*), and all other bonds by the LINCS method (*43*). In simulations without ammonium, K^+^ and Cl^-^ ions were added to neutralize the system and obtain a bulk ionic concentration of 250 mM. In simulations with ammonium, K^+^ was replaced by NH_4_^+^. The parameters for NH_4_^+^ and NH_3_ (umbrella sampling simulations) were adapted from Nygaard *et al*. (*23*). As the protonation pattern of the twin-His motif has been found to play a role in the hydration of the protein (*24*), two different tautomeric states of the twin-His motif were systematically probed in the simulations. The tautomeric state in which H168 is protonated on its N_δ_ and H318 is protonated on its N_ε_ atom is referred to as ‘DE’, while ‘ED’ terms the twin-His configuration where H168 is protonated on N_ε_ and H318 is protonated on N_δ_ (Fig. S7).

After a steepest descent energy minimization, the system was equilibrated by six consecutive equilibration steps using position restraints on heavy atoms of 1000 kJ/mol.nm^2^. The first three equilibration steps were conducted in an NVT ensemble, applying a Berendsen thermostat (*44*) to keep the temperature at 310K. The subsequent steps were conducted under an NPT ensemble, using a Berendsen barostat (*44*) to keep the pressure at 1 bar. Production molecular dynamics simulations were carried out using a v-rescale thermostat (*45*) with a time constant of 0.2 ps, and a Berendsen barostat with semi-isotropic coupling. A timestep of 2 fs was used throughout the simulations.

In a subset of simulations, we aimed to test the effect of membrane voltage on the internal hydration of AmtB using CompEL. For the CompEL simulations (*46*), the system was duplicated along the z-axis, perpendicular to the membrane surface. To focus on the physiologically relevant voltage gradient in *E. coli*, i.e. a negative potential on the inside of the cell of magnitude −140 to −170 mV (*47*), an antiparallel orientation of the two trimers in the double bilayers was used (*46*). The final double system consisted of a rectangular box of 13×13×20nm. For the CompEL simulations, 1000 positively charged (either NH_4_^+^ or K^+^) and 1000 negatively charged ions (Cl^-^) were added to the system, then the system was neutralized, and the desired ion imbalance established.

The Umbrella Sampling (US) Potential-of-Mean-Force (PMF) calculations (*48*) were set up as described previously by Hub et al. (*16*). A snapshot was taken from the simulation of the single bilayer system with the twin-His motif in the DE protonation state and both the CWW and PWW occupied. The pore coordinates were obtained using the software HOLE (*49*), removing the solvent and mutating F215 to alanine during the HOLE run only. Starting coordinates for each umbrella window were generated by placing NH_3_ in the central x-y coordinate of the pore defined by HOLE at positions every 0.5 Å in the z coordinate. Solvent molecules within 2 Å of the ammonia’s N atom were removed. Minimisation and equilibration were then re-performed as described above. Unless otherwise stated, position restraints were used for all water oxygen atoms in the CWW with a 200 kJ/mol.nm^2^ force constant; while the TIP3 molecules within the lower water wire were not restrained. For the US the N atom of ammonia was position-restrained with a force constant of 1000 kJ/mol.nm^2^ on the z axis and a 400 kJ/mol.nm^2^ cylindrical flat-bottomed potential with a radius of 5 Å in the x-y plane, as described earlier by Hub et al. (*16*). For some US window simulations, the ammonia z-axis restraints were increased and the time step reduced in the equilibration to relax steric clashes between sidechains and ammonia. After equilibration, US simulations were run for 10ns, using the parameters described above (*39*) and removing the initial 2 ns for further equilibration. The PMF profiles were generated with the GROMACS implementation of the weighted histogram analysis method (WHAM) with the periodic implementation (*50*). Further US simulations were performed to as needed to improve sampling in regions of the profile that were not sufficiently sampled. The Bayesian bootstrap method was performed with 200 runs to calculate the standard deviation of the PMF.

### Free energy calculations for proton translocation

The free energies for proton translocation were evaluated by protonating the water molecules at different sites along the periplasmic and cytoplasmic water wires. Electrostatic effects in proteins are often treated more effectively using semi-macroscopic models which can overcome the convergence problems of more rigorous microscopic models. Here we used the semi-macroscopic protein dipole/Langevin dipole approach of Warshel and coworkers in the linear response approximation version (PDLD/S-LRA) (*51, 52*). Positions of the water molecules in the PWW and CWW were obtained from the corresponding MD snapshots (Fig. 1). All PDLD/S-LRA p*K*_a_ calculations were performed using the automated procedure in the MOLARIS simulations package (*53*) in combination with the ENZYMIX force field. The simulation included the use of the surface-constrained all atom solvent model (SCAAS) (*54*) and the local reaction field (LRF) long-range treatment of electrostatics. At each site, 20 configurations for the charged and uncharged state were generated. The obtained p*K*_a_ values were then converted to free energies for proton translocation.

## Acknowledgments

Special thanks to Prof. Iain Hunter (Strathclyde Institute of Pharmacy and Biomedical Sciences) for invaluable discussions and help during this project. We also thank Pascale Van Vooren for technical support and Thomas P. Jahn for sharing the *triple-mepΔ trk,1,2Δ* yeast strain.

## Authors contributions

AJ and UZ suggested the research topic. AB, AJ, AMM, AP, CMI, ET, GW, GDM, GT, MBa, MBo, UZ and AJ designed, performed the research and analysed the data. UZ and AJ wrote the manuscript. All authors contributed to the discussion of the results and editing the manuscript. The authors have no competing interests.

## Supplementary Materials

### Supplementary figures

**Fig. S1:**
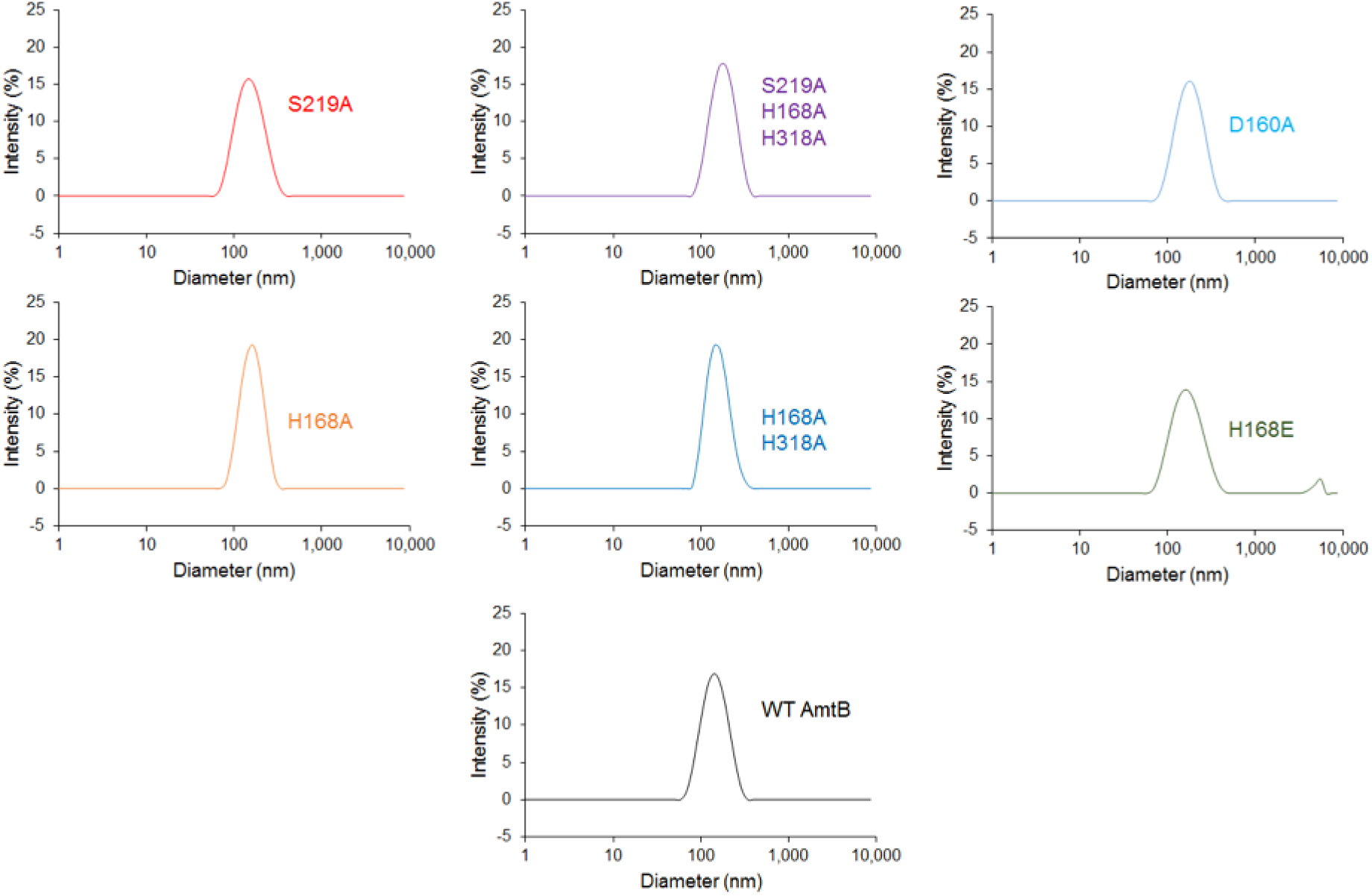
Size distribution of the proteoliposomes containing wild-type and variants of AmtB. Dynamic light scattering was used to determine the number-weighted distribution of liposome sizes in the detection reagent. The distribution was monodisperse, with a mean diameter of 110 nm.

**Fig. S2:**
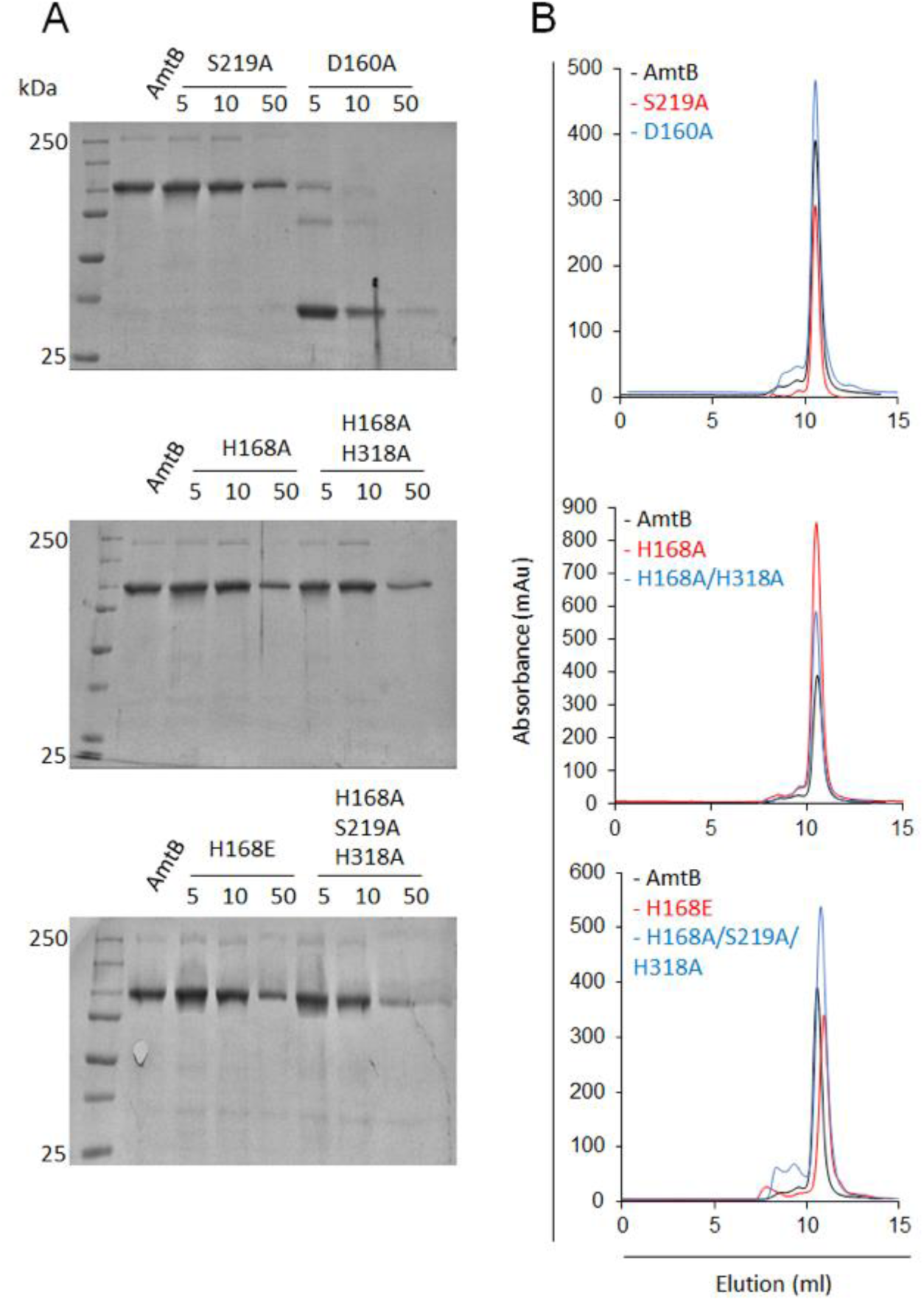
Quatification of AmtB variants in proteoliposomes. **(A)** SDS-PAGE Coomassie Blue– stained gel of the proteoliposome containing AmtB variants. 15 µL of proteoliposomes (5mg/ml) containing each variant at a LPR of 5:1, 10:1, or 50:1 (w/w), or wild-type AmtB at LPR 10 (WT-AmtB) were analysed on 10% SDS-PAGE gels. **(B)** Gel filtration trace (Superdex 200 10/300 increase) of wild-type AmtB and variants after solubilization of the proteoliposome in 2% DDM. All of the AmtB variants elute at ∼ 11.5 ml.

**Fig. S3:**
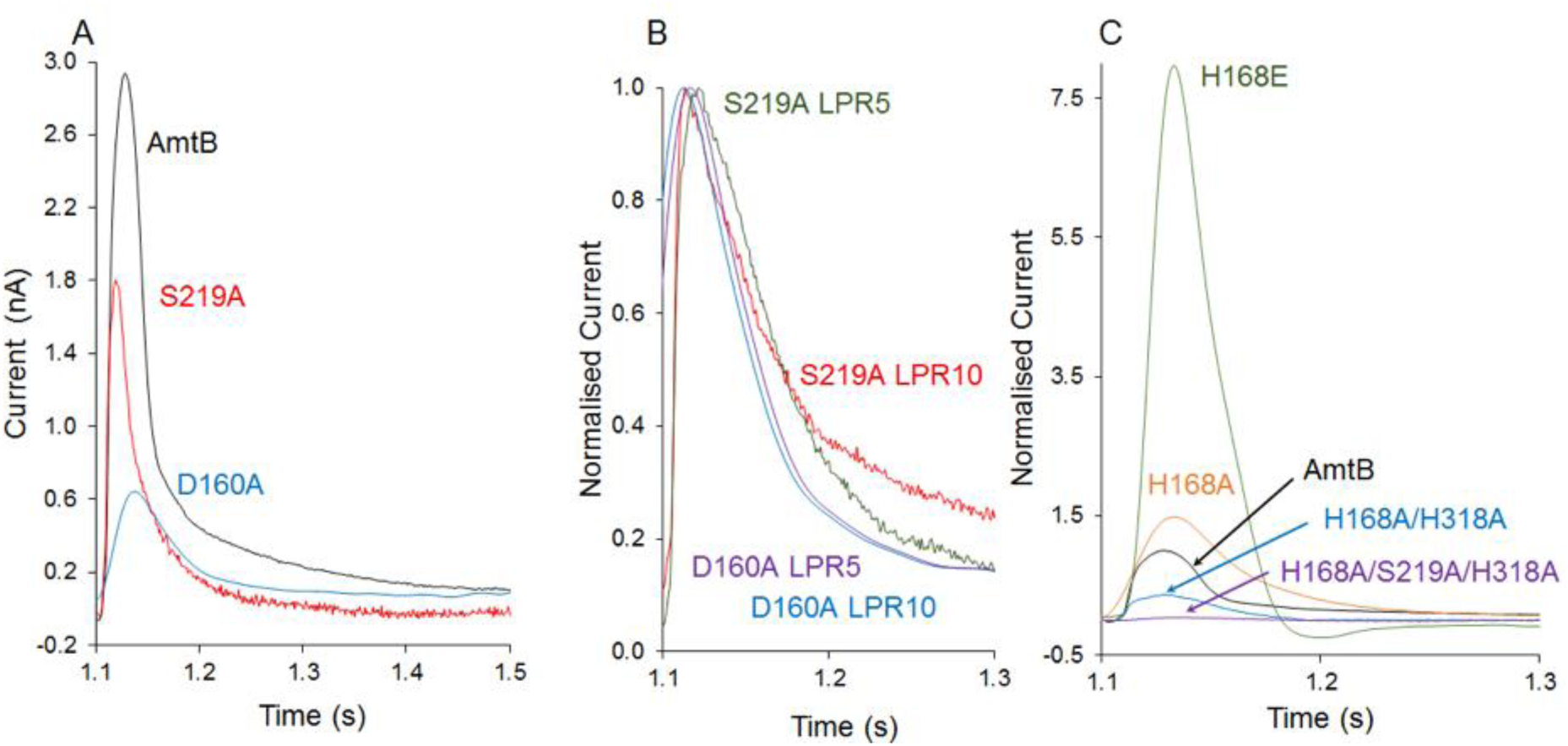
Characterization of the activity and specificity of AmtB variants. **(A)** SSME analysis. Transient current measured after a 200 mM pulse of ammonium in proteoliposomes containing AmtB (black), AmtB^S219A^ (red), and AmtB^D160A^ (blue) at LPR 10. **(B)** Normalized traces following a 200 mM ammonium pulse in proteoliposomes containing AmtB^S219A^ at LPR 10 (red), AmtB^S219A^ at LPR 5 (green), AmtB^D160A^ at LPR 10 (blue), and AmtB^D160A^ at LPR 5 (purple). **(C)** SSME analysis. Transient current measured after a 200 mM ammonium pulse in proteoliposomes containing AmtB (black), AmtB^H168AH318A^ (Blue), AmtB^S219AH168A318A^ (purple), AmtB^H168E^ (green), and AmtB^H168A^ (orange).

**Fig. S4:**
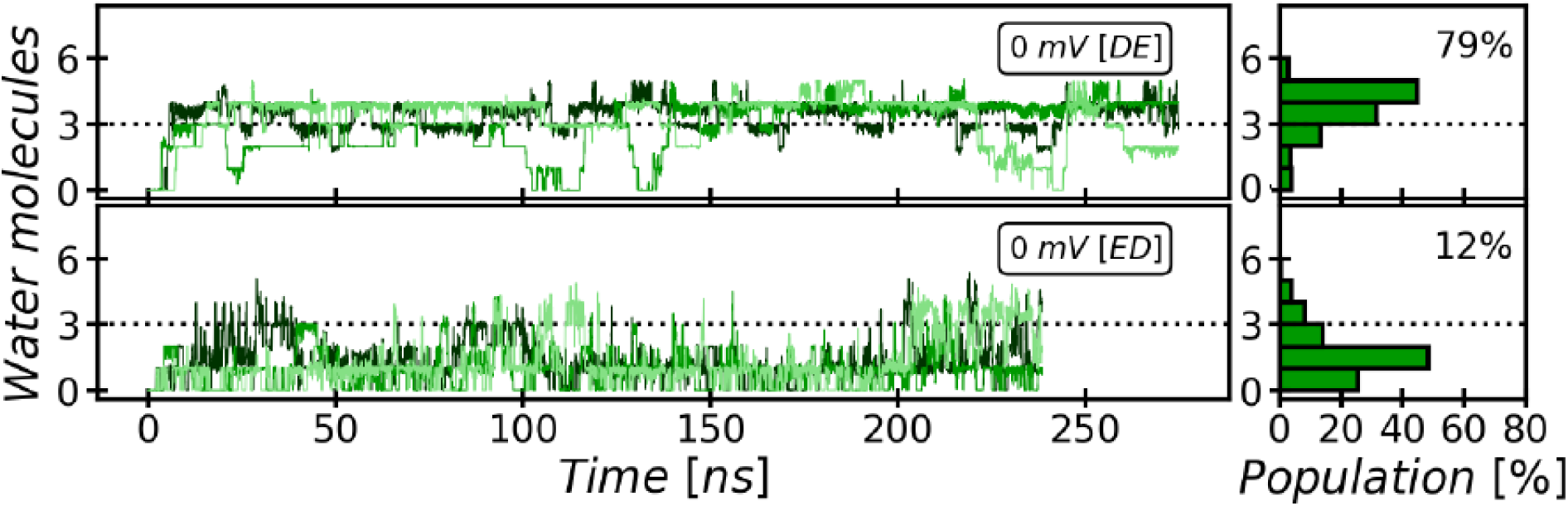
Evolution and occupancy of the Cytoplasmic Water Wire (CWW). The cytoplasmic water wire develops in an early stage of the simulations, as has been previously observed (*28*). The CWW is particularly stabilized in the DE twin-His tautomeric state, where the occupancy is 79% on average with at least three water molecules over the course of our simulations.

**Fig. S5:**
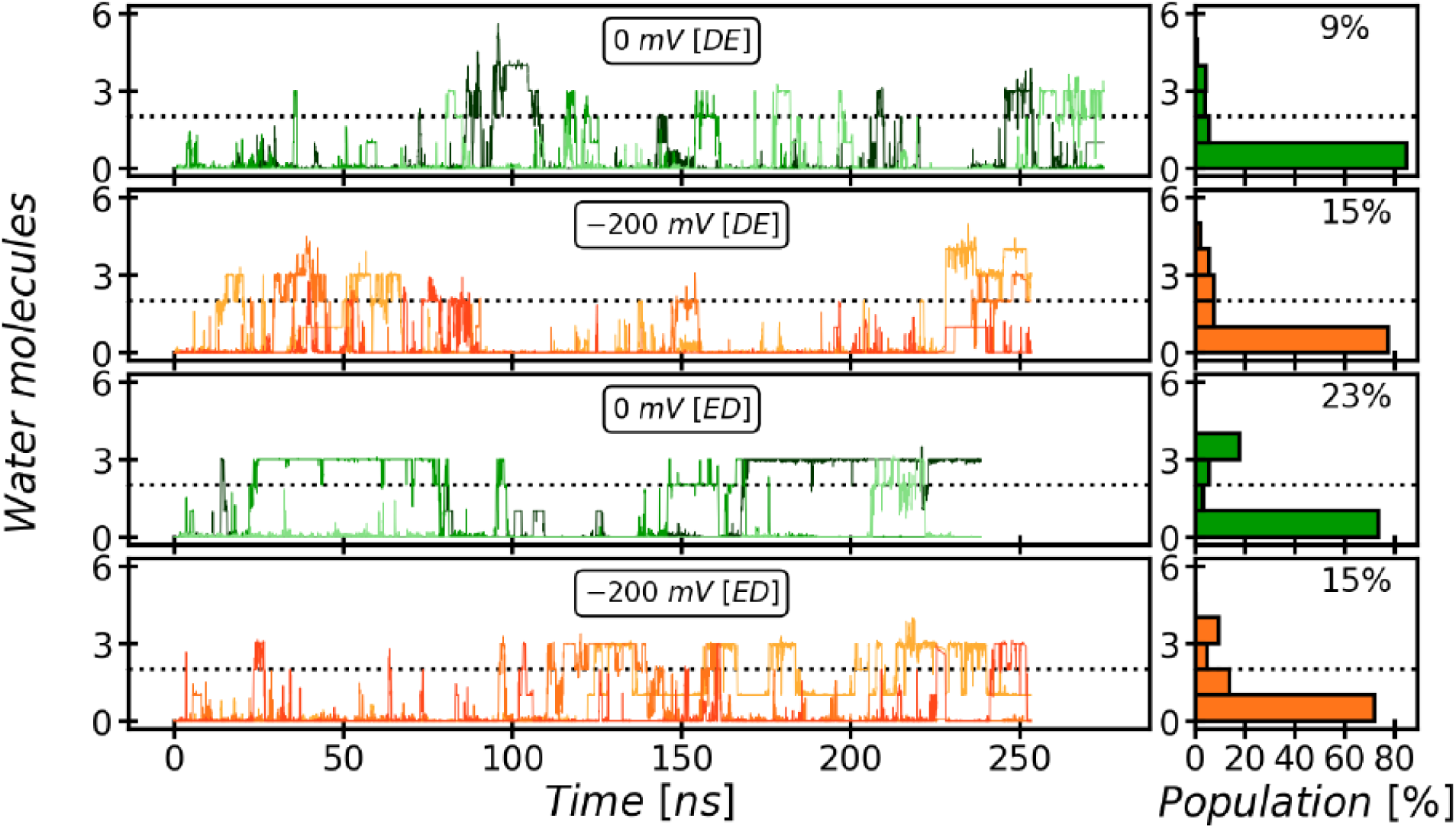
Evolution and occupancy of the Periplasmic Water Wire (PWW). The periplasmic water wire evolves on a slower time scale than the CWW and shows greater fluctuations. It is stabilized by the ED twin-His tautomeric state, where 23% occupancy of the PWW with three water molecules or more is observed. Stably occupied states are seen after ∼25 ns simulated time. Since earlier simulations studying the internal hydration of AmtB were shorter (*28*), this may explain why the PWW has not been described previously. A significant influence of membrane voltage of a magnitude that is biologically relevant for the cytoplasmic membrane of *E. coli* is not seen.

**Fig. S6:**
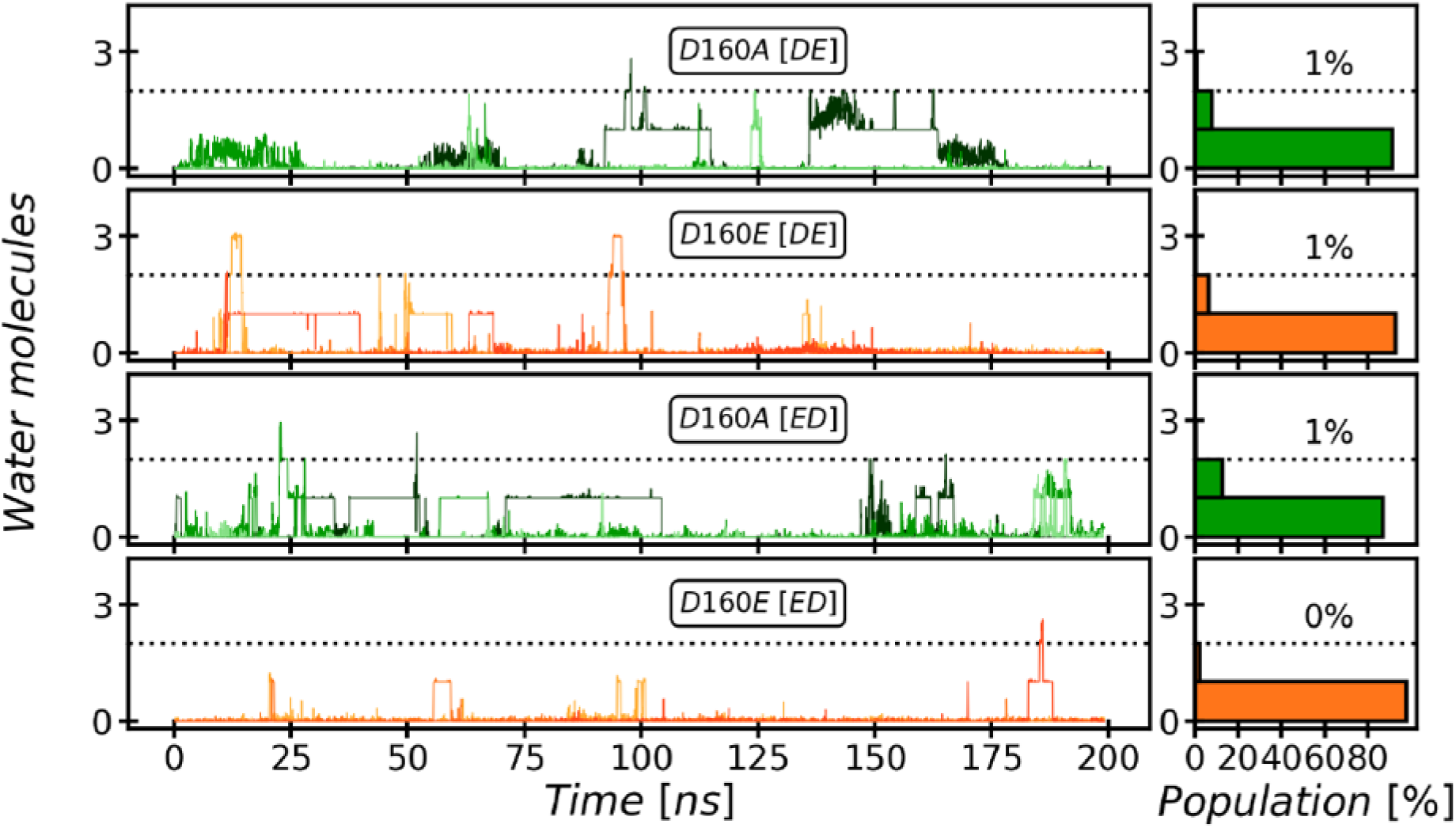
The Periplasmic Water Wire (PWW) in the D160A and D160E variants. We observe no significant occupancy of the PWW above the threshold of at least three water molecules in the D160A and D160E AmtB variants, irrespective of the tautomeric protonation states of H168 and H318.

**Fig. S7:**
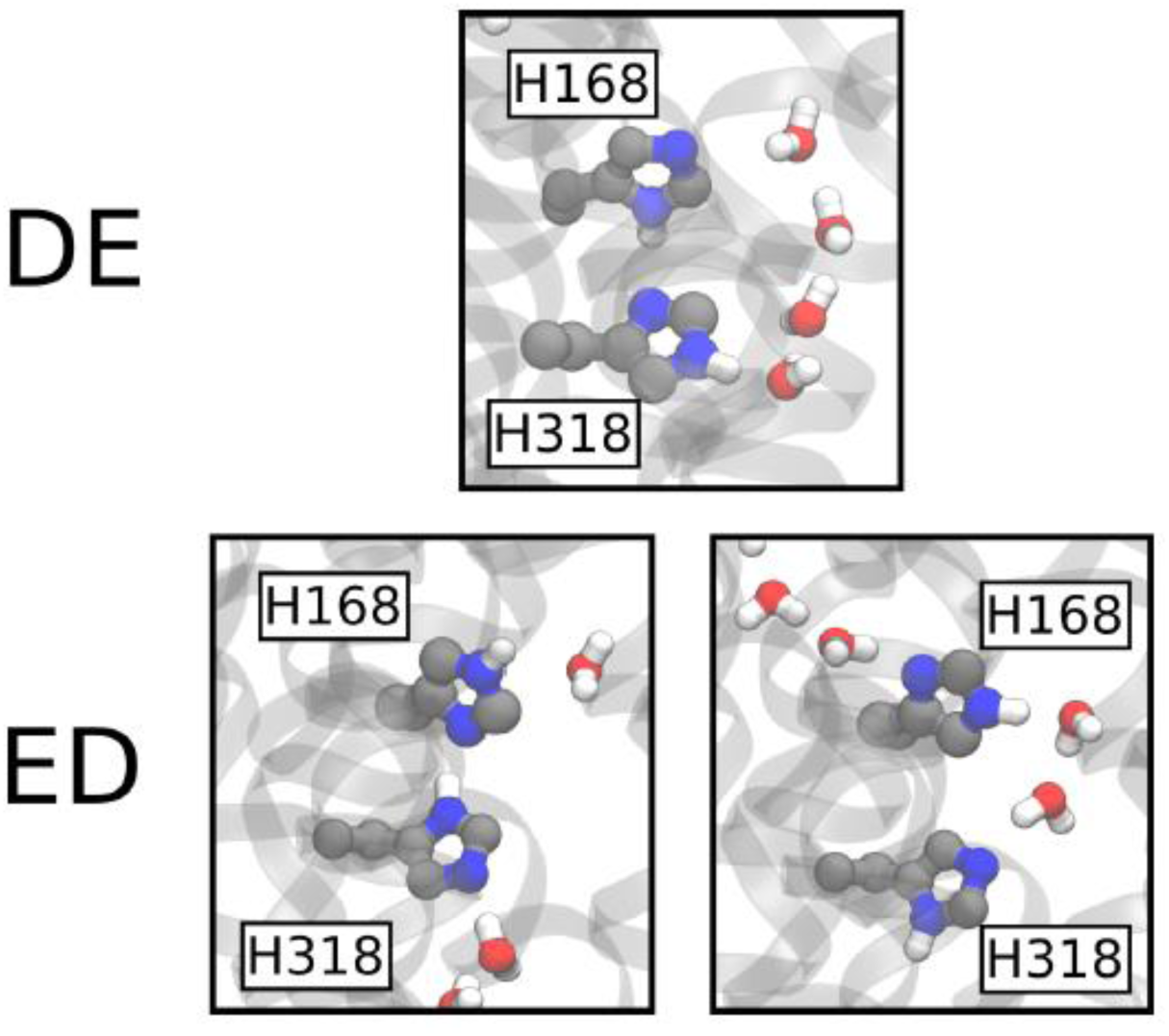
The DE and ED twin-His motif configurations. Nomenclature as in Lamoureux *et al*. (*28*). The top panel shows the configuration of H168 and H318 observed in the DE simulations, with the protonated Nδ atom of H168 pointing towards the Nδ atom of H318. The bottom panel shows the two distinct configurations observed in the ED simulations; on the left is the equivalent of the situation observed in DE, while on the right the H168 sidechain flips and its Nδ atom contacts the most proximal molecule of the PWW.

**Fig. S8:**
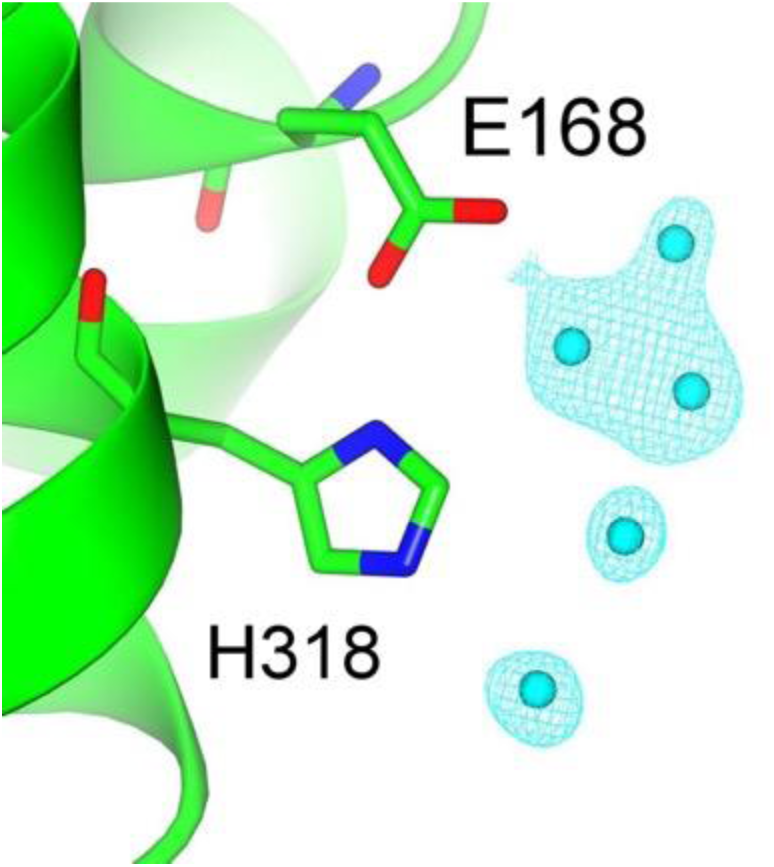
Modelled water molecules and electron density (cyan) around E168 and H318 in AmtB variant H168E. Crystallographic data and analysis taken from Javelle *et al.* (*21*)

**Fig. S9:**
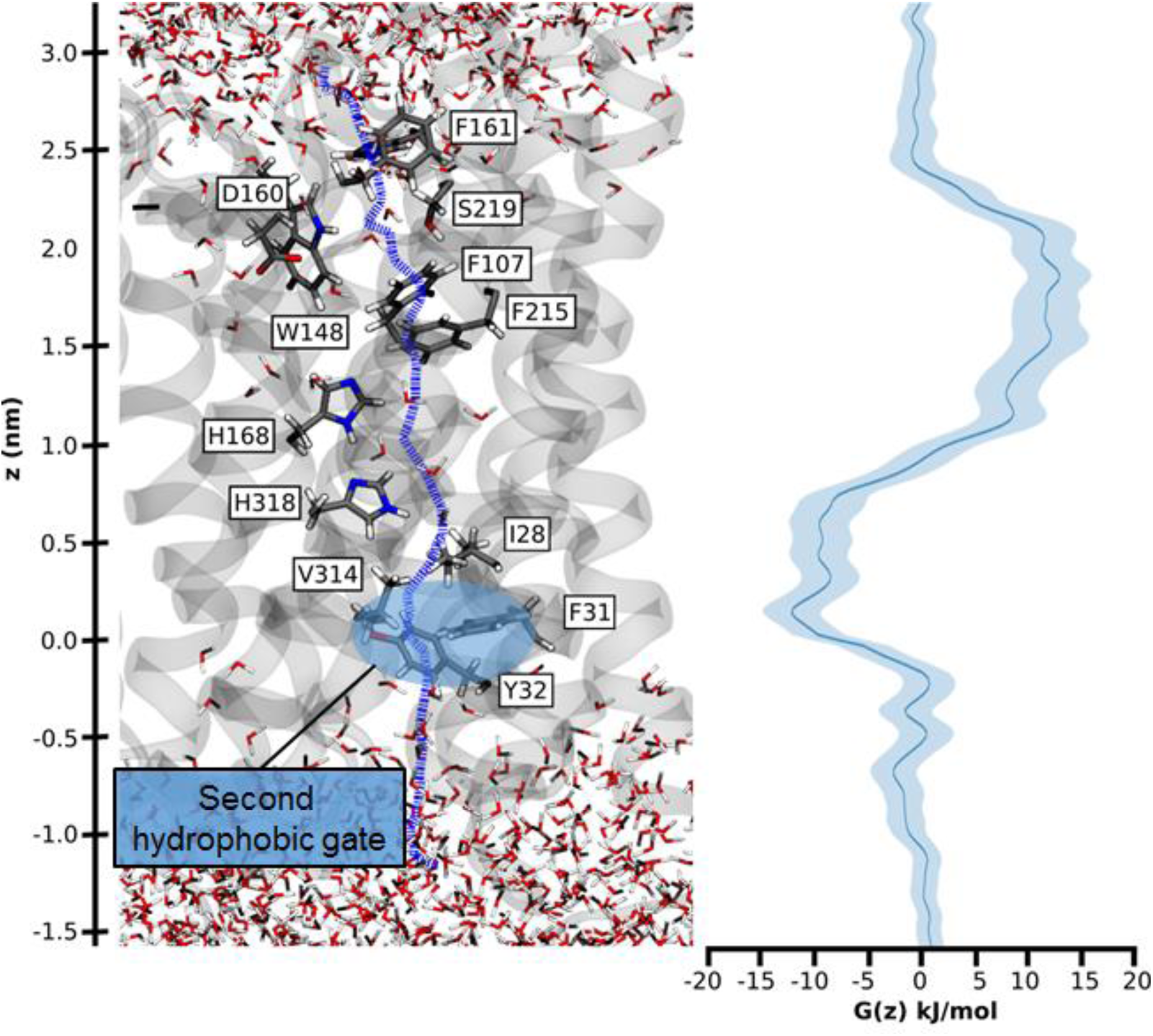
Hydrophobic pathway and energetics for NH_**3**_ translocation in AmtB. We probed an optimal pathway for NH_3_ transfer during our PMF calculations (left, purple dash trajectory) in the presence of both the PWW and CWW. The software HOLE (*49*) was used to determine the most likely transfer route. The pathway from the periplasm to the cytoplasm traverses the hydrophobic gate region (F107 and F215), crosses the cavity next to the twin-His motif (H168 and H318) occupied by the CWW, and continues across a second hydrophobic region (I28, V314, F31, Y32) before entering the cytoplasm. The energetics of transfer on this pathway (right, blue curve) show that NH_3_ translocation experiences only a moderate energy barrier during traversal of the periplasmic hydrophobic gate region (∼10 kJ/mol, near F107 and F215). By contrast, a shallow binding site is observed near the cytoplasmic hydrophobic gate (∼-5 kJ/mol). The increased residence time of NH_3_ within this energy minimum suggests that reprotonation to NH_4_^+^, caused by the cytoplasmic pH, likely occurs near this site, plausibly at its cytoplasmic exit.

### Supplementary tables

**Table S1:**
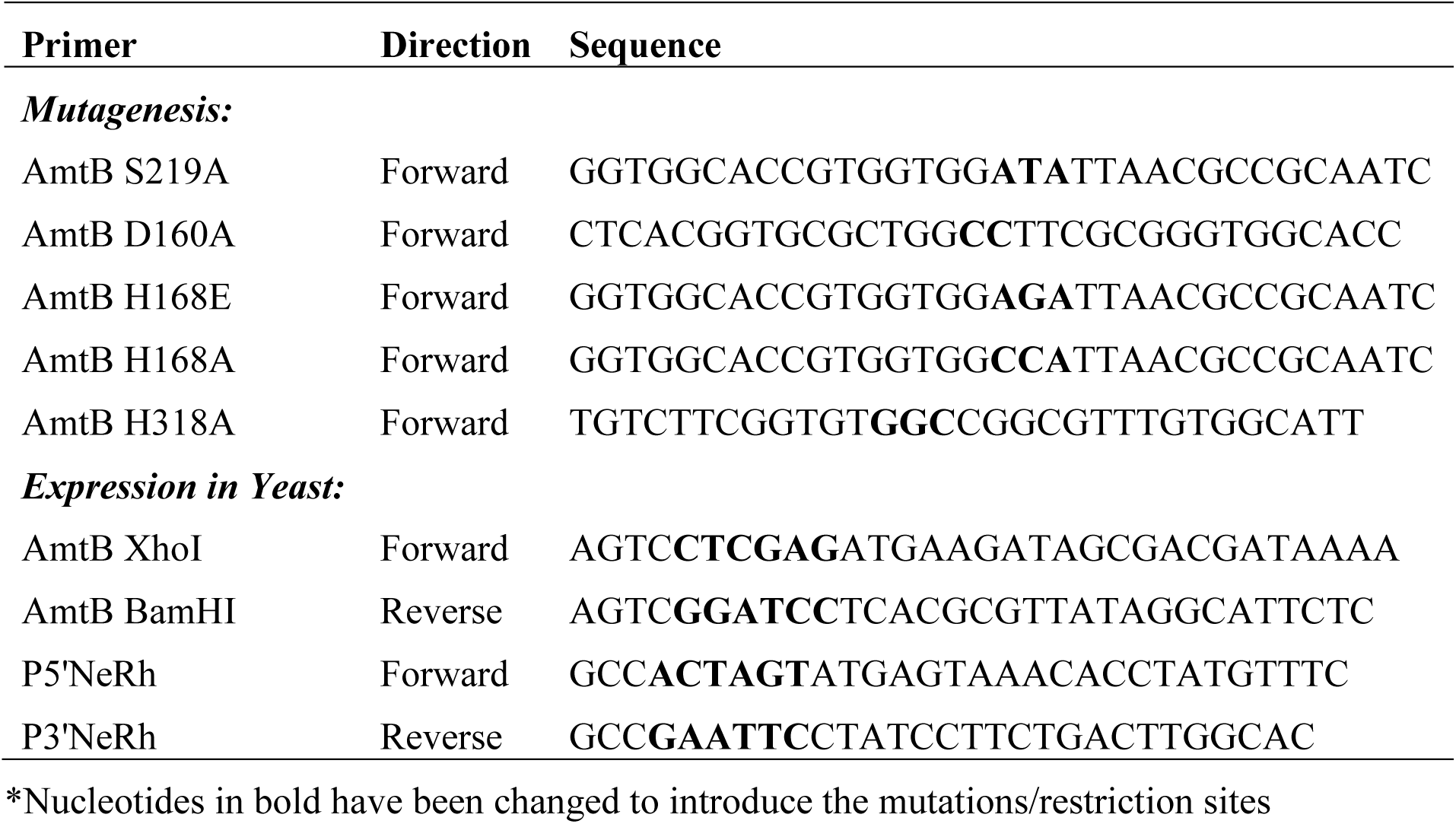
Primer use in this study*

**Table S2:** Plasmids use in this study

| Plasmid | Description | Reference |
| --- | --- | --- |
| pET22b (+) |  | Novagen |
| pDR195 | High copy yeast expression vector | (55) |
| pAD7 | pESV2-Rh50(His)<sub>6</sub> | (15) |
| p426MET25 |  | (56) |
| pZheng | pET22b-AmtB(His)<sub>6</sub> | (34) |
| pAJ2024 | pET22b-AmtB(His)<sub>6</sub><sup>S219A</sup> | (57) |
| pGDM2 | pET22b-AmtB(His)<sub>6</sub><sup>H168AH318A</sup> | This study |
| pGDM4 | pET22b-AmtB(His)<sub>6</sub><sup>D160A</sup> | This study |
| pGDM6 | pET22b-AmtB(His)<sub>6</sub><sup>S219AH168AH318A</sup> | This study |
| pGW1 | pET22b-AmtB(His)<sub>6</sub><sup>H168E</sup> | This study |
| pGW2 | pET22b-AmtB(His)<sub>6</sub><sup>H168A</sup> | This study |
| pGDM7 | pDR195-AmtB<sup>S219A</sup> | This study |
| pGDM9 | pDR195-AmtB<sup>D160A</sup> | This study |
| pGDM12 | pDR195-AmtB<sup>H168AH318A</sup> | This study |
| pGDM13 | pDR195-AmtB<sup>S219AH168AH318A</sup> | This study |
| pGW6 | pDR195-AmtB<sup>H168E</sup> | This study |
| pGW7 | pDR195-AmtB<sup>H168A</sup> | This study |

**Table S3:**
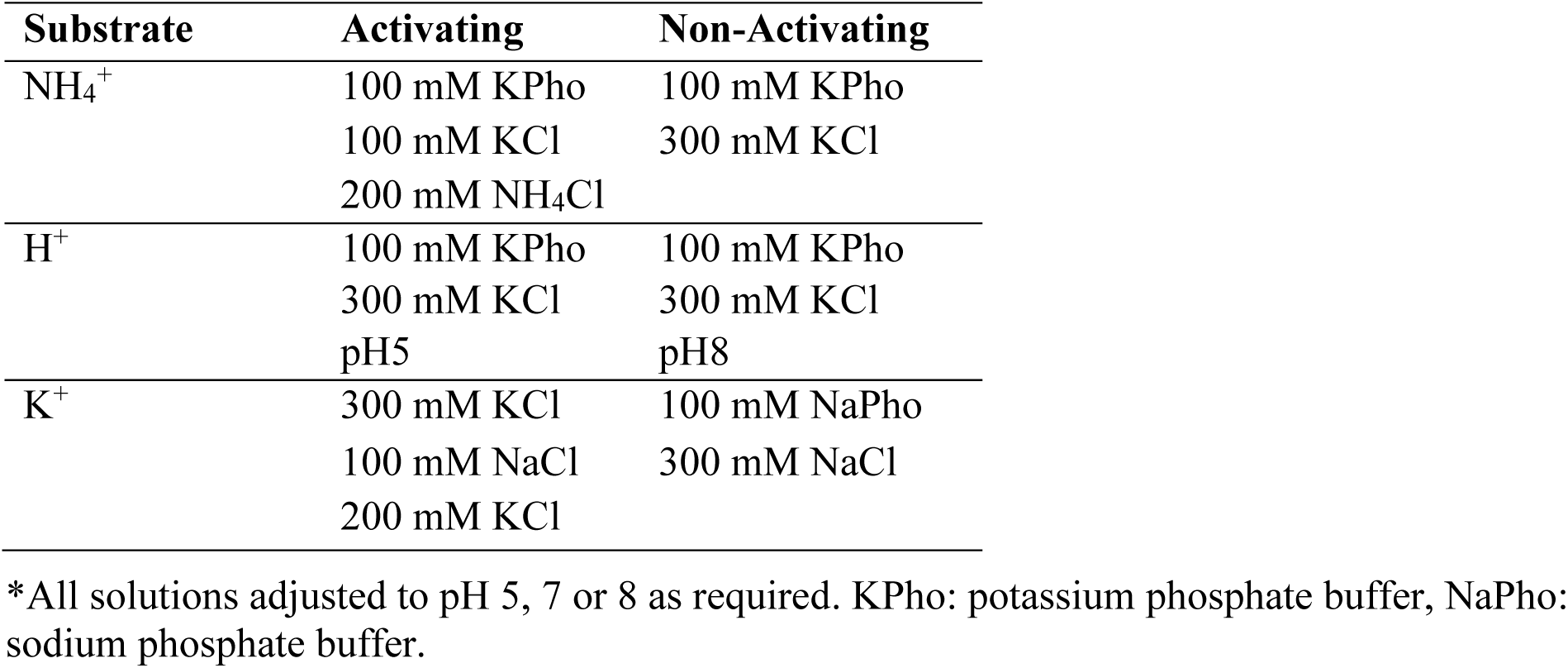
Solid-Supported Membrane Electrophysiology Solutions*

**Table S4:** Decay time constants (s^-1^) of transient currents triggered after an ammonium or potassium pulse of 200 mM in proteoliposomes containing AmtB at various LPR*.

| **Variant** | **NH<sub>4</sub><sup>+</sup>** |  | **K<sup>+</sup>** |  |
| --- | --- | --- | --- | --- |
|  | **LPR 10** | **LPR 5** | **LPR 10** | **LPR 5** |
| **AmtB-WT** | 13.4 ± 1.5 | 18.7±1.0 | NC | NC |
| **S219A** | 15.0 ± 0.6 | 22.9±1.4 | NC | NC |
| **D160A** | 21.6 ± 1.2 | 24.3 ± 1.5 | NC | NC |
| **H168A H318A** | 31.5 ± 1.1 | 31.2 ± 3.6 | NC | NC |
| **S219A H168A H318A** | NC | NC | NC | NC |
| **H168A** | 28.3 ± 1.5 | 38.0 ± 1.0 | 2.7 ± 0.5 | 5.2 ± 1.0 |
| **H168E** | 30.4 ± 1.3 | 40.2 ± 1.0 | NC | NC |
| **NeRh50** | 24.0 ± 1.7 | 39.0 ± 3.6 | NC | NC |

\*NC: No transient current recorded

**Table S5:**
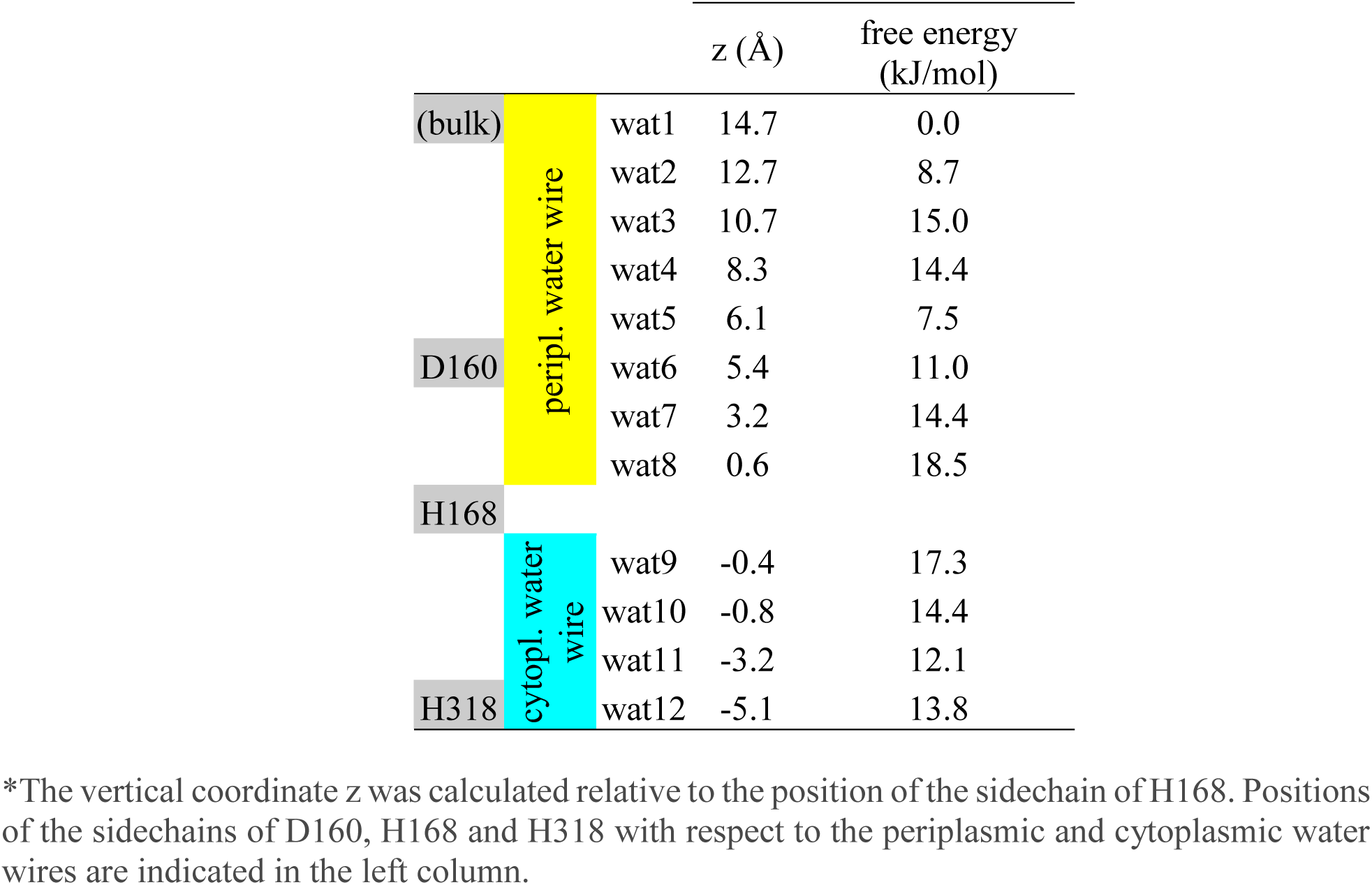
Free energies for proton translocation through the cytoplasmic and periplasmic water wires and neighboring water molecules (bulk)*.

